# Regulation of the Rheb GTPase via GATOR1 Complex

**DOI:** 10.64898/2025.12.20.695697

**Authors:** Aditi Prabhakar, William B. Mair

## Abstract

mTORC1 coordinates cellular growth and metabolism by integrating inputs from both amino acids and growth factors, and its activation requires two upstream branches involving the Rag GTPases and the Rheb GTPase. These branches are regulated by distinct GAP complexes: GATOR1 (Depdc5-Nprl2-Nprl3) inhibits RagA/B, and TSC (TSC1-TSC2-TBC1D7) inhibits Rheb. Despite the prevailing view that these pathways converge only at mTORC1 itself, several observations suggest upstream crosstalk. This gap is especially striking in organisms like *C. elegans* and *S. cerevisiae* that lack the TSC complex yet maintain fully responsive mTORC1 signaling. How these inputs are dynamically coordinated under complex physiological conditions and in organisms lacking the key components remain unknown. We performed unbiased quantitative proteomics in *C. elegans* and identified the GATOR1 complex as a previously unrecognized RHEB-1 (*C. elegans* ortholog of Rheb) interactor. Through biochemical validation in human cells, we show that nucleotide-free Rheb associates with the Nprl2–Nprl3 subunits of GATOR1, whereas GTP-bound or membrane-detached Rheb mutants fail to bind. Nutrient stress, but not direct pharmacologic inhibition of mTORC1, robustly induced this interaction. In TSC2-null cells, where Rheb is constitutively GTP-loaded, Rheb-Nprl2/3 binding was strongly diminished and was restored by expressing the nucleotide-free Rheb^S20N^ mutant, demonstrating that Rheb’s nucleotide state governs this interaction. Pulldown assays confirmed that the Nprl2/3 heterodimer is sufficient for binding nucleotide-free Rheb. Structural modeling using AlphaFold3 consistently positioned Rheb at a conserved site on Nprl3 distinct from the RagA/B GAP-active surface of Nprl2, supporting a non-catalytic mode of association. Together, these findings identify a conserved, nutrient-regulated physical interaction between Rheb and the Nprl2/3 subunits of GATOR1, revealing a previously unrecognized point of convergence between the growth factor and amino acid branches of the mTORC1 pathway. This model provides a direct molecular link between the Rag and Rheb branches, furthering our understanding of how nutrient stress fine-tunes mTORC1 signaling.

## INTRODUCTION

Cells must coordinate their growth and metabolism with extracellular cues such as growth factors and nutrient availability. The mechanistic target of rapamycin complex 1 (mTORC1) is a serine/threonine kinase that serves as a central hub in this process and integrates diverse signals such as growth factors, nutrient availability (amino acids and glucose), cellular energy levels, and stress cues^1^. When active, mTORC1 promotes anabolic processes including protein, lipid, and nucleotide synthesis, while suppressing catabolic processes like autophagy, thus coupling the decision to grow with sufficient resources (**Fig. 1A**). Deregulation of mTORC1 signaling is implicated in diseases ranging from metabolic disorders such as insulin resistance and obesity to cancer, underscoring its fundamental role in cellular homeostasis^1^.

**Figure 1.**
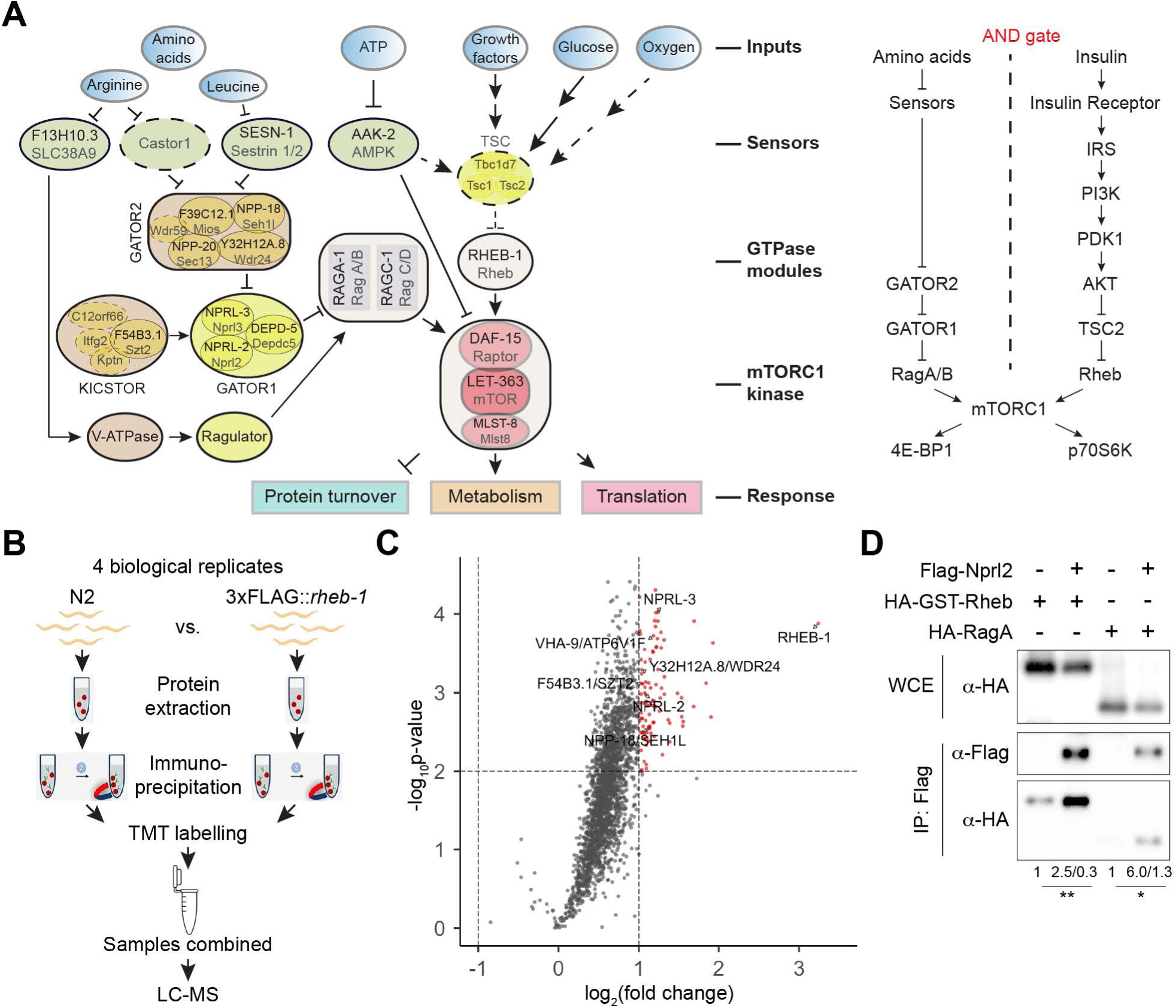
Rheb/RHEB-1 physically associates with subunits of the nutrient-sensing GATOR1 complex. **(A)** The mTORC1 pathway. Left, schematic of the integrated signaling network showing how multiple cues converge at either Rheb or RAG GTPases to activate the mTORC1 kinase complex. Color code: blue, inputs; green, sensors that directly bind the inputs; grey, GTPases; yellow, GAPs (TSC and GATOR1) and GEF (Ragulator) for the GTPases; brown, other scaffolding and regulatory complexes; red, the mTORC1 kinase complex. Black font, *C. elegans* orthologs; grey font, mammalian orthologs. Solid outlines, proteins conserved in *C. elegans*; dotted outlines, proteins absent in *C. elegans*. Arrows indicate activating signals; perpendicular lines indicate inhibitory signals. Right, simplified genetic pathway illustrating the AND-gate model of mTORC1 activation, which requires concurrent input from growth-factor (Rheb) and amino-acid (RAG) signaling branches. **(B)** Schematic of the *C. elegans* proteomic workflow used to identify endogenous RHEB-1 interactors. Immunoprecipitation-mass spectrometry (IP-MS) was performed on endogenously tagged 3×FLAG::*rheb-1* animals compared with untagged wild-type controls. **(C)** Volcano plot showing proteins enriched in RHEB-1 immunoprecipitates relative to untagged wild type across four biological replicates. Dotted lines indicate statistical cut-offs: a two-fold enrichment and p < 0.01. Red dots denote hits that pass both thresholds. **(D)** Co-immunoprecipitation (CO-IP) of FLAG-Nprl2 with HA-RagA (positive control) and HA-GST-Rheb in HEK293T cells under basal conditions. Values below the blots represent mean fold-enrichment and standard error of mean from six independent replicates, each normalized to its control IP, which was set to 1. *, p < 0.05; **, p < 0.01.

mTORC1 activation at the lysosome is coordinated by two key upstream signaling branches: one governed by the small GTPase Rheb^2,3^ and the other by the Rag GTPases^4^. Rheb directly activates mTORC1 when in its GTP-bound state, while the Rag GTPases mediate nutrient-dependent localization of mTORC1 to the lysosomal membrane. In the presence of insulin or other growth factors, receptor tyrosine kinase activation triggers the PI3K-Akt pathway, which phosphorylates and inhibits the tuberous sclerosis complex (TSC1-TSC2-TBC1D7)^5^. The TSC complex acts as a GTPase-activating protein (GAP) for Rheb, maintaining Rheb in an inactive GDP-bound state and thereby suppressing mTORC1 activity^6,7^. When Akt inactivates TSC2, this brake is relieved, allowing Rheb to accumulate in its GTP-bound active form. GTP-loaded Rheb localizes to lysosomal membranes and directly binds to mTORC1, inducing a conformational change that activates mTORC1’s kinase activity^2^. Rheb is essential for mTORC1 activation by all upstream cues. Concurrently, amino acid availability is sensed through a parallel pathway centered on the Rag family of small GTPases. The Rag GTPases (heterodimers of RagA or RagB with RagC or RagD) reside on the cytosolic face of lysosomes, and in amino acid-replete conditions they adopt an active nucleotide state (RagA/B^GTP^-RagC/D^GDP^)^4^. Active Rag heterodimers bind Raptor (an mTORC1 subunit), thereby recruiting the mTORC1 complex to lysosomal membranes. This relocation is crucial because it brings mTORC1 into proximity with active Rheb, enabling Rheb to trigger mTORC1 kinase activity. In contrast, under amino acid starvation, the Rag GTPases are driven into an inactive nucleotide state (RagA/B^GDP^-RagC/D^GTP^), preventing mTORC1 lysosomal localization and thus, blocking its activation. A key regulator of this nutrient-sensing branch is the GATOR1 complex (a heterotrimer of Depdc5, Nprl2, and Nprl3), which functions as the GAP for RagA/B^8^.

These two branches act as a two-input coincidence detector (analogous to an “AND-gate”); Rheb-dependent signaling and Rag-dependent localization must both be engaged to achieve full mTORC1 activation^1^. This paradigm ensures that mTORC1 is optimally active only when both growth factor signals and nutrient signals are present, thereby preventing inappropriate anabolic activity during nutrient deprivation or growth factor withdrawal. According to prevailing models, these pathways converge only at the level of mTORC1 itself, with no direct crosstalk upstream of the complex. Nonetheless, hints of functional interaction between the nutrient-sensing and growth factor pathways have existed for some time. Notably, amino acid starvation not only inactivates Rag GTPases but also triggers lysosomal recruitment of the TSC complex, bringing the Rheb GAP into proximity of Rheb^9,10^. Conversely, loss of GATOR1 renders mTORC1 signaling insensitive to growth factor withdrawal, such that mTORC1 remains aberrantly active upon PI3K-Akt inhibition even when amino acid inputs are absent, suggesting the amino acid branch can influence the efficacy of growth factor withdrawal-mediated downregulation^11^. These observations imply that the nutrient and growth factor inputs may intersect upstream of mTORC1.

A striking and unresolved issue is that several model organisms, including *C. elegans* and *S. cerevisiae*, lack core components of the canonical AND gate (most notably the entire TSC complex)^12,13^, yet retain upstream PI3K-Akt signaling, Rheb, Rag GTPases, and a fully functional, nutrient- and growth factor-responsive mTORC1 pathway^1,14^ (**Fig. 1A**). How these organisms achieve coordinated regulation of Rheb and Rag inputs in the absence of TSC remains unknown. Thus far, no molecular node of integration has been identified that could reconcile how mTORC1 remains responsive to complex physiological conditions, where nutrient and hormonal inputs dynamically intersect, as well as how organisms missing key AND-gate components accomplish this integration. To address these gaps, we turned to an unbiased approach of quantitative proteomics in *C. elegans* to identify novel regulators of RHEB-1 (*C. elegans* ortholog of Rheb).

Here, we report the identification of a direct point of convergence between the two mTORC1 regulatory branches – at the level of Rheb itself. Through a combination of quantitative proteomics in *C. elegans*, *in vitro* biochemical validation in human cells, and AlphaFold3^15^ structural modeling, we identified a previously unrecognized physical interaction between Rheb and the GATOR1 complex. Specifically, co-immunoprecipitation studies revealed that nucleotide-free Rheb (apo-Rheb) binds to the Nprl2-Nprl3 heterodimer of GATOR1. This unexpected Rheb-GATOR1 association is stimulated by upstream nutrient stress such as amino acid deprivation, whereas direct pharmacologic inhibition of mTORC1 activity did not induce this interaction. Mechanistically, the interaction was specific to Rheb’s nucleotide state, only inactive, apo-Rheb (nucleotide-free) bound GATOR1. A mutant Rheb locked in a nucleotide-empty state (Rheb^S20N^) co-precipitated strongly with Nprl2, whereas GTP-bound (Rheb^Q64L^) or mislocalized (Rheb^C181S^) mutants showed little to no binding. Consistent with this, in cells lacking TSC2 (where Rheb is constitutively GTP-loaded) the Rheb-GATOR1 interaction was greatly diminished but could be restored by introducing the nucleotide-free Rheb mutant. Through pulldown assays, we further mapped this interaction to the Nprl2/3 subunits of GATOR1, indicating that the Nprl2-Nprl3 heterodimer alone is sufficient to bind Rheb in its apo state. AlphaFold3 modeling placed the Rheb-binding site on Nprl3, well outside the canonical GAP domain on Nprl2^16^, suggesting a non-catalytic, sequestration-type mode of interaction. Together, these findings uncover a direct, nutrient-regulated linkage between the amino acid-sensing machinery and the Rheb branch of mTORC1 signaling.

## RESULTS

### Rheb physically associates with the nutrient-sensing complex GATOR1

To identify conserved regulators of Rheb, we performed an unbiased proteomic screen in *C. elegans* (**Fig. 1B-C**). Because *C. elegans* lacks the TSC complex^12,13^ that acts as the canonical GAP for Rheb in mammals, this system provides a simplified context to uncover upstream or parallel regulators of the Rheb branch of mTORC1. Tandem-mass-tag (TMT) quantitative IP-mass spectrometry^17^ was performed on animals expressing endogenously tagged 3xFLAG::*rheb-1* compared with untagged wild type controls (N2) (**Fig. 1B, Fig. S1**) across four biological replicates. The analysis revealed consistent enrichment of several components belonging to the amino acid-sensing arm of the mTORC1 pathway (**Fig. 1C**). Among the most significantly enriched interactors were the *C. elegans* orthologs of Nprl2 (NPRL-2) and Nprl3 (NPRL-3), core subunits of the GATOR1 complex; Wdr24 (Y32H12A.8) and Seh1l (NPP-18), components of GATOR2 complex; Szt2 (F54B3.1), the sole *C. elegans* ortholog of the KICSTOR complex; and ATP6V1F (VHA-9), a component of the lysosomal V-ATPase. This suggests that RHEB-1 can associate with several members of the amino-acid-sensing machinery, providing a potential physical point of crosstalk that could underlie coordination between the Rheb and Rag branches.

To determine whether this interaction is conserved in mammalian cells, we co-expressed FLAG-Nprl2 with HA-GST-Rheb or HA-RagA (used as the positive control) in HEK293T cells (**Fig. 1D**). As expected, FLAG-Nprl2 robustly co-immunoprecipitated HA-RagA (6-fold enrichment over control IPs). FLAG-Nprl2 also recovered HA-GST-Rheb (2.5-fold enrichment over control IPs), indicating that Rheb can physically associate with Nprl2 in mammalian cells. Together, these data identify GATOR1 subunits as a previously unrecognized Rheb-binding complex, revealing a conserved physical connection between the growth-factor-sensing Rheb branch and the amino-acid-sensing GATOR1 branch of mTORC1 regulation.

To begin defining the molecular mechanism underlying this association, we turned to the human orthologs expressed in HEK293T cells, which provide a tractable system for dissecting regulatory inputs under controlled nutrient and signaling conditions.

### Nutrient availability regulates the Rheb-Nprl2 interaction

Having established that Rheb physically associates with the GATOR1 subunit Nprl2, we next asked whether this interaction is regulated by cellular nutrient status. Because mTORC1 integrates both growth-factor and nutrient inputs, we tested conditions that selectively inhibit each upstream branch to determine whether either cue alters Rheb–Nprl2 association.

We first tested whether growth-factor deprivation influences the interaction. Cells expressing FLAG-Nprl2 and HA-GST-Rheb were exposed to serum starvation for 16 h, which reduces growth-factor signaling through the PI3K-AKT-TSC axis, and led to a modest decrease in mTORC1 activity as measured by phospho-p70S6K levels (**Fig. S2**). Under this condition, RagA and Rheb co-immunoprecipitated strongly with Nprl2 compared with basal medium, indicating growth-factor deprivation influences the Rheb-Nprl2 interaction (**Fig. S2**).

We next tested whether amino-acid availability similarly modulates the interaction. Amino-acid starvation independently suppresses mTORC1 signaling via GATOR1-dependent inhibition of the Rag GTPases, which prevents mTORC1 recruitment to lysosomes^8,18^. Cells expressing FLAG-Nprl2 and HA-GST-Rheb were incubated in amino-acid-free medium for four hours. Loss of amino acids markedly decreased phospho-p70S6K levels, confirming effective inhibition of mTORC1 (**Fig. 2A, S2**). In these samples, the Nprl2-RagA association remained intact, validating the immunoprecipitation conditions. Amino-acid deprivation also increased the amount of HA-GST-Rheb recovered with FLAG-Nprl2 (**Fig. 2A-B, S2)**. These findings demonstrate that amino-acid starvation also enhances Rheb-Nprl2 association in human cells. Although the Rheb-Nprl2 interaction increased under both serum and amino acid starvation relative to basal conditions, neither insulin nor amino acid refeeding clearly disengaged Rheb from Nprl2. This likely reflects the incomplete recovery of mTORC1 activity under our re-stimulation conditions and is further supported by the parallel failure of RagA to dissociate from Nprl2 during re-stimulation (**Fig. S2**). Together, these results demonstrate that nutrient deprivation enhances Rheb-Nprl2 association in human cells, while re-stimulation conditions used here were insufficient to reverse this effect.

**Figure 2.**
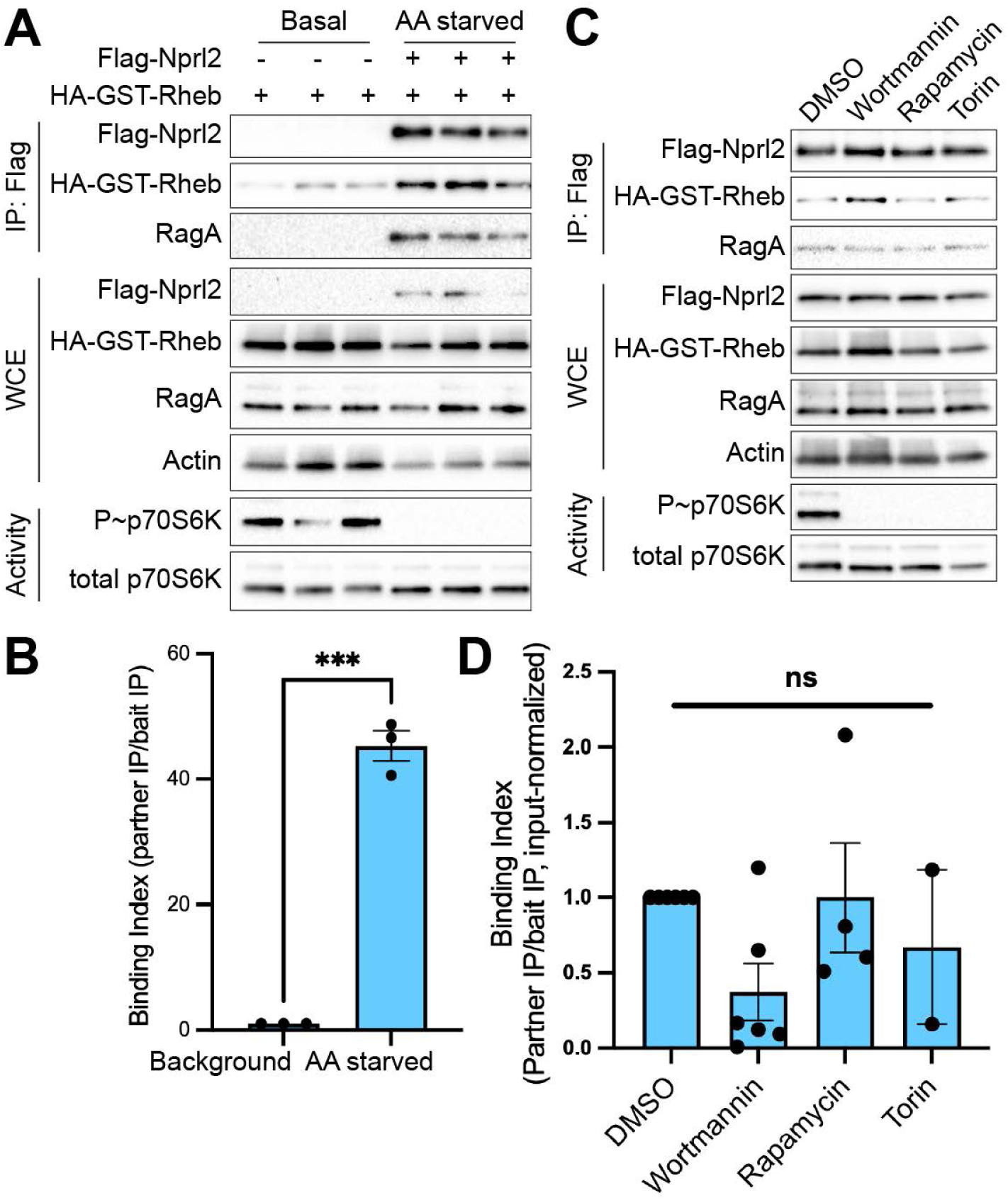
Rheb-GATOR1 interaction is a nutrient-responsive, upstream event. **(A)** HEK293T cells expressing indicated plasmids were incubated in amino acid-free media for 4 h before CO-IP analyses. Total expression shown in whole-cell extracts (WCE). mTORC1 activity was monitored by phospho-p70S6K levels in whole-cell extracts. **(B)** Quantification of panel A. Bars, mean; error bars, standard error of mean (SEM). Individual dots represent independent biological replicates. **(C)** HEK293T cells expressing FLAG-Nprl2 and HA-GST-Rheb were treated for 1 h with DMSO (vehicle), Wortmannin (PI3K inhibitor), Rapamycin (allosteric mTORC1 inhibitor), or Torin1 (ATP-competitive mTOR inhibitor) before CO-IP analyses. **(D)** Quantification of panel C done as in panel B. One-way ANOVA was used for statistical significance. *** p < 0.001; ns, not significant.

Because Rheb-Nprl2 association consistently coincided with conditions of reduced mTORC1 activity, we next asked whether this interaction simply reflects a general consequence of mTORC1 inhibition or arises from upstream regulatory events. To distinguish between these possibilities, we used pharmacological inhibitors that directly block mTORC1 kinase activity without altering its nutrient or growth-factor inputs. Treatment with Wortmannin (a PI3K inhibitor that also directly inhibits the mTOR kinase^19,20^), Rapamycin (an allosteric mTORC1 inhibitor^21–23^), or Torin1 (an ATP-competitive mTOR inhibitor^24^) each robustly suppressed mTORC1 signaling, as confirmed by the loss of phospho-p70S6K (**Fig. 2C**). However, none of these inhibitors increased the association of either RagA or Rheb with Nprl2 compared with vehicle-treated control. Quantification across independent replicates showed no significant difference in Nprl2-bound Rheb levels relative to baseline (**Fig. 2D**).

Together, these results demonstrate that the Rheb-Nprl2 interaction is selectively induced by nutrient deprivation and is not a downstream consequence of mTORC1 inhibition, indicating that the regulation occurs upstream of mTORC1 kinase activity.

### Rheb-Nprl2 association preferentially involves the inactive form of Rheb

Because the activity of small GTPases is tightly governed by their nucleotide-bound state, we next asked whether the association between Rheb and Nprl2 depends on Rheb’s nucleotide status. As a member of the Ras superfamily, Rheb possesses a conserved GTP-binding domain containing switch I and switch II regions that mediate conformational changes upon GTP or GDP binding, as well as a C-terminal CaaX motif that anchors Rheb to membranes via farnesylation^25–29^ (**Fig. 3A**). These conserved features have enabled the generation of well-characterized alleles that lock Rheb in distinct conformations: Rheb^S20N^, which impairs nucleotide binding and mimics a nucleotide-free or an inactive state^30^; Rheb^Q64L^, which is GTP-locked due to defective TSC binding^31^; and Rheb^C181S^, which disrupts farnesylation and thus membrane association^25^. We co-expressed HA-Nprl2 with FLAG-tagged Rheb alleles in HEK293T cells and assessed their ability to co-immunoprecipitate (**Fig. 3B-C**). Phospho-p70S6K analysis confirmed that the Rheb alleles behaved as expected, with Rheb^S20N^ exhibiting reduced mTORC1 activity and Rheb^Q64L^ showing enhanced activation of p70S6K kinase relative to wild-type Rheb^6,30^. Consistent with previous reports, loss of farnesylation and membrane localization in Rheb^C181S^ did not abolish Rheb’s ability to stimulate S6K phosphorylation^30^ (**Fig. 3B**). We found that relative to the wild-type Rheb, the S20N allele exhibited a pronounced increase in Nprl2 association. Notably, FLAG-Rheb^S20N^ expression was markedly lower than that of other alleles, yet the co-immunoprecipitation signal remained the strongest among all variants. The S20N mutant is known to be unstable and expressed at very low levels (Gerta Hoxhaj, personal communication). The construct was independently validated by plasmid sequencing and by confirming reduced mTORC1 activity. In contrast to S20N mutant, the GTP-locked Q64L mutant and the cytosolic C181S mutant displayed only weak association with Nprl2. Together, these data demonstrate that the inactive or nucleotide-free conformation of Rheb preferentially associates with Nprl2, suggesting that Nprl2 engages Rheb when it is in an inactive or nucleotide-transition state.

**Figure 3.**
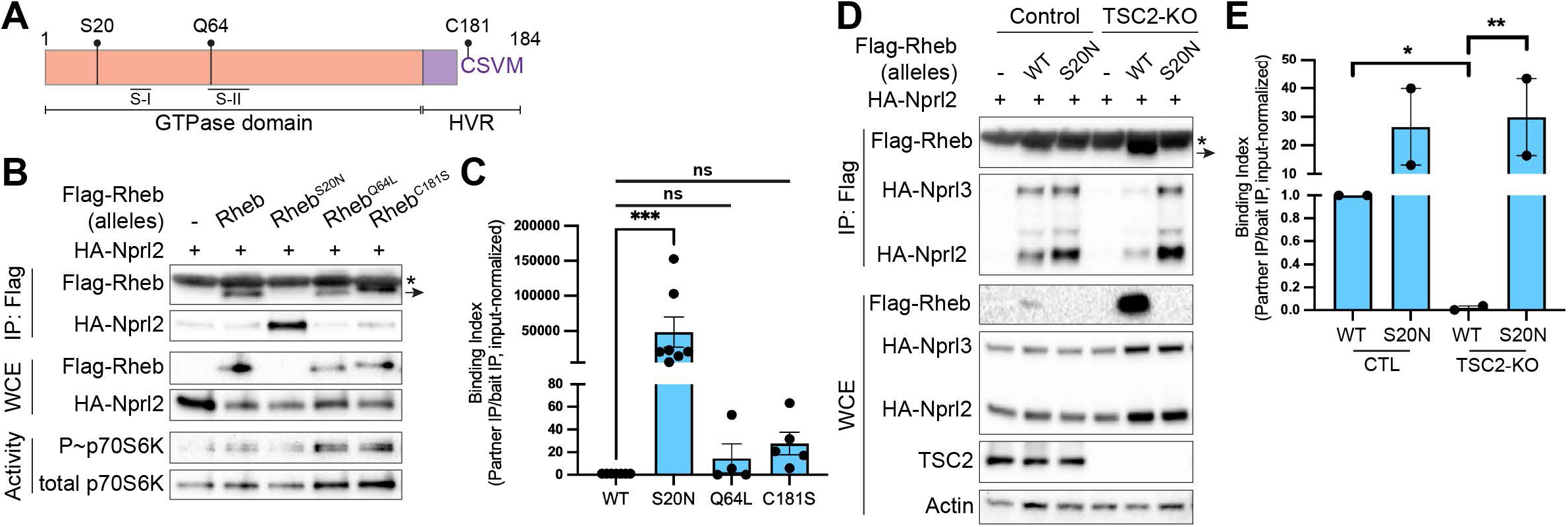
The Rheb-Nprl2 interaction depends on the nucleotide state of Rheb. **(A)** Schematic of Rheb functional domains and the alleles used to test nucleotide- and localization-dependent effects on Rheb-Nprl2 binding. Rheb contains conserved switch I (S-I) and switch II (S-II) regions in its N-terminal GTPase domain that mediate conformational changes upon nucleotide exchange and a C-terminal hypervariable region (HVR) ending in CaaX motif required for farnesylation and membrane localization. Relative position of critical residues is shown. **(B)** HEK293T cells were co-transfected with HA-Nprl2 and indicated FLAG-tagged Rheb variants for CO-IP analysis. *, light chain signal from FLAG-magnetic beads; arrow, FLAG-Rheb signal. FLAG-Rheb^S20N^ expression is markedly reduced relative to other alleles. **(C)** Quantification of panel B same as in Fig. 2. One-way ANOVA was used for statistical significance. **(D)** HEK293T cells co-expressing indicated FLAG-tagged Rheb variants with HA-Nprl2 and HA-Nprl3 in control and TSC2-knockout cells for CO-IP analyses. **(E)** Quantification of panel D same as in Fig. 2. Unpaired t-test with Welch’s correction was used for statistical significance between indicated pairs. *, p < 0.05; **, p < 0.01; ***, p < 0.001; ns, not significant.

To further assess this model in a cellular context, we examined Rheb-Nprl2 binding in a TSC2-null background, where Rheb accumulates in its active GTP-bound form due to loss of the TSC GAP activity^30^. In the TSC2-knockout cells, the association of wild-type Rheb with Nprl2 was markedly diminished relative to control cells, consistent with the predominance of the GTP-bound form of Rheb when TSC2 GAP activity is lost. (**Fig. 3D-E**). Strikingly, expression of Rheb^S20N^ in the TSC2-null background restored strong co-immunoprecipitation with Nprl2, further confirming that the inactive or nucleotide-free state of Rheb is the favored configuration for Nprl2 engagement.

In this experimental setup, we also co-expressed HA-Nprl3 to determine whether the interaction extends to other GATOR1 subunits. Nprl3 co-immunoprecipitated with Rheb in a pattern similar to Nprl2, strong binding with the S20N allele and weak binding with wild-type Rheb in TSC2-null cells (**Fig. 3D**), indicating that either the entire GATOR1 complex or the Nprl2-Nprl3 heterodimer (henceforth, Nprl2/3) can associate with inactive Rheb. Collectively, these findings establish that the Rheb-Nprl2/3 interaction is nucleotide-state-dependent and that Nprl2/3 preferentially engages the inactive or nucleotide-free form of Rheb.

### Rheb interacts with GATOR1 through a structurally consistent but transient interface predicted by AlphaFold3

To understand the structural nature and regulatory determinants of the Rheb-GATOR1 interaction, we performed *in silico* modeling using AlphaFold3^15^ across multiple conditions. To further dissect whether the full GATOR1 complex or its Nprl2/3 subcomplex can interact with inactive Rheb, we systematically modeled the Rheb-GATOR1 complex using human orthologs under diverse conditions, including the presence or absence of Depdc5, different Rheb alleles, including wild-type (WT), the GTP-locked Q64L mutant, and the nucleotide-free S20N mutant, as well as with multiple random seeds to account for model variability. Despite differences in complex composition, Rheb mutation, or seed number, the predicted binding site of Rheb remained remarkably consistent across all models. In ribbon-structure visualizations, Rheb docked onto the same interface at Nprl3, occupying a conserved surface distinct from the GAP-active groove and the canonical RagA/B-binding site (**Fig. 4A**, top panels, blue sphere marks the GAP domain which interacts with RagA^16^). This robust spatial consistency across models, even in the absence of Depdc5 (**Fig. 4A**, bottom panels) supports the idea that Rheb can associate with either the full GATOR1 complex or the Nprl2/3 subcomplex alone.

**Figure 4.**
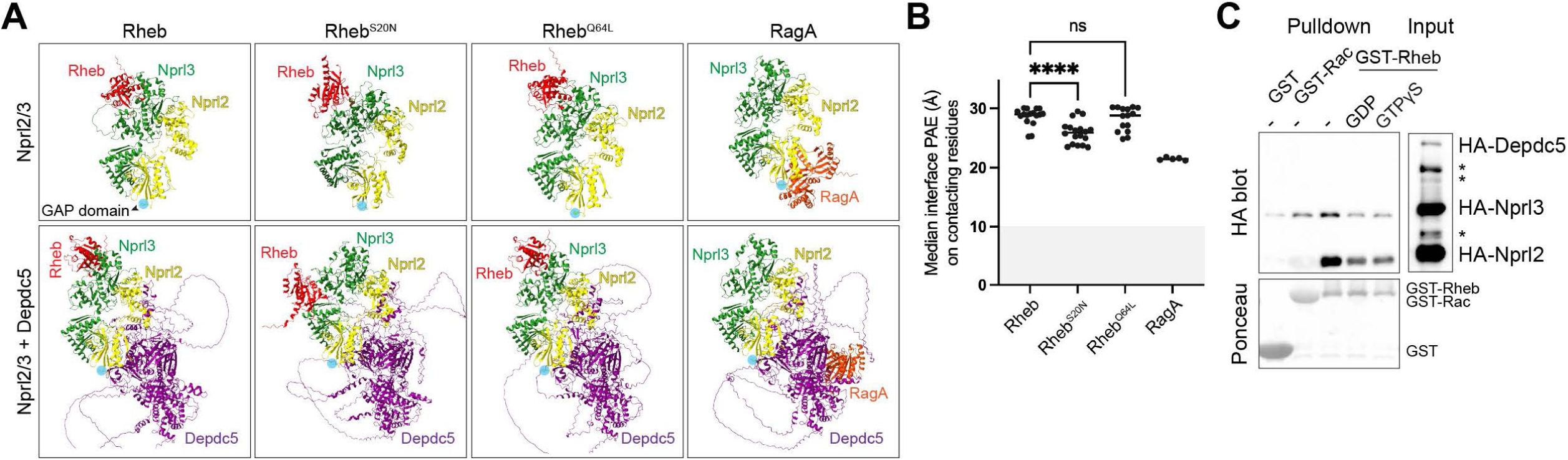
Nucleotide-free Rheb predicted to bind a conserved site on the Nprl2/3 heterodimer. **(A)** AlphaFold3-predicted ribbon structures of Rheb or RagA modeled with the human Nprl2/3 heterodimer (top) or the full GATOR1 complex (bottom, with Depdc5 included). Each complex was modeled with Rheb^WT^, Rheb^S20N^ (nucleotide-free mimic), or Rheb^Q64L^ (GTP-bound mimic). Color scheme: Rheb (red), RagA (orange), Nprl2 (yellow), Nprl3 (green), Depdc5 (purple). Blue circle, GAP domain within Nprl2. **(B)** Median predicted alignment error (PAE) for contacting residues at the Rheb-Nprl3 interface, averaged across all models for each condition, with and without Depdc5. RagA-Nprl2 is shown as a positive control. Gray shaded region denotes the gold-standard for high-confidence threshold for rigid, obligate interactions (PAE < 10 Å). Individual dots, each model’s median interface PAE value. ****p < 0.0001; ns, not significant. One-way ANOVA with Dunnett’s post hoc test was used for significance. **(C)** HEK293T cells overexpressing HA-tagged GATOR1 components were subjected to amino acid starvation (4 h), and lysates were incubated with bacterially purified GST-Rheb, GST-Rac, or GST alone, preloaded with no nucleotide (–), GDP (500 μM) or GTPγS (500 μM). Pulldowns were analyzed by immunoblots for HA epitope; GST fusion proteins were visualized by Ponceau S staining. Input shows relative expression of tagged GATOR1 subunits. Asterisks indicate non-specific background bands.

AlphaFold3 provides multiple quantitative metrics that together inform the confidence and accuracy of predicted complex assemblies. These include predicted Local Distance Difference Test (pLDDT) scores, which estimate the confidence in the position of each residue within its own chain; predicted TM-score (pTM), which reflects the overall confidence in the global fold of the complex; and inter-chain predicted TM-score (ipTM), which specifically evaluates the confidence in inter-chain interfaces. Additionally, the Predicted Aligned Error (PAE) matrix estimates the expected error in positioning any residue relative to another, serving as a critical tool for assessing the precision of domain or subunit orientation. Collectively, these scores help differentiate between well-folded chains, confidently predicted interfaces, and regions with flexibility or uncertainty.

When examining the AlphaFold3-predicted structures of Rheb-Nprl2/3 and RagA-Nprl2/3 complexes, we observed that the overall model confidence was modest across all conditions. Both the global fold confidence (pTM) and inter-subunit interface confidence (ipTM) were consistently low, and this trend was reflected in the residue-level pLDDT scores. Notably, none of the models for either Rheb or RagA bound to the Nprl2/3 heterodimer alone reached the typical confidence thresholds (ipTM > 0.6, pTM > 0.5) required to infer a high-confidence interface. This pattern was observed even for the RagA-Nprl2 interface, a biochemically validated interaction, suggesting that AlphaFold may report low confidence when modeling transient or context-specific interactions (**Fig. S3**, top panels). Strikingly, inclusion of the Depdc5 subunit in our AlphaFold3 models markedly increased the confidence of the predicted Rheb-Nprl2/3 and RagA-Nprl2/3 complex structures (**Fig. S3**, bottom panels). Because AlphaFold and experimental studies reliably stabilize regions rendered ordered only in the context of full complexes^32,33^, adding Depdc5 supplies the structural context needed for higher-confidence prediction of the Rheb-Nprl2/3 and RagA-Nprl2/3 assemblies.

Although the binding interface remained structurally consistent, we observed meaningful differences in AlphaFold’s predicted confidence for this interaction based on Rheb’s nucleotide state. We quantified the PAE at the Rheb-Nprl3 contact for each Rheb variant. Rheb^S20N^’s interface PAE (∼25) is modestly but consistently lower than WT/Q64L (∼28) (**Fig. 4B**), suggesting that AlphaFold interprets that nucleotide-free Rheb is the most favored configuration for engaging at this site. Furthermore, quantification of valid contact models (defined as models containing Rheb-Nprl3 residue pairs ≤8 Å apart) revealed that Rheb^S20N^ formed a contact with Nprl3 in 18 out of 20 models, compared to 17 for wild-type and 15 for the GTP-locked Q64L, suggesting a failure to engage the interface tightly in some Q64L models. These findings further reinforce that Rheb’s nucleotide state governs its ability to interact with GATOR1.

It is worth noting that all absolute PAE values were relatively high. To assess whether high PAE values at this interface necessarily indicate a non-real binding or might reflect a real but transient interaction, we used the RagA-Nprl2 interface as a biological benchmark. The GATOR1 complex is a validated GAP for RagA/B, and its engagement with RagA has been structurally and biochemically well established^16,34^. Yet, AlphaFold3 predictions for the RagA-Nprl2/3 complex similarly yielded high PAE values (**Fig. 4B**) and low ipTM scores (**Fig. S3**, ∼0.4), consistent with a flexible interface that can adopt multiple binding modes. Thus, AlphaFold also treats RagA-Nprl2, a nutrient-gated contact as weak or transient. These data support the interpretation that a high PAE may reflect a legitimate but transient or low-affinity interaction^35^, fitting with the known biology of both RagA and Rheb engagement with GATOR1.

To experimentally validate these *in silico* observations, we performed GST-pulldown assays to directly test whether nucleotide-free Rheb preferentially engages the GATOR1 complex over its GDP-bound counterpart. We purified recombinant GST-tagged Rheb preloaded with either no nucleotide (–), GDP, or the non-hydrolyzable GTP analog GTPγS, and incubated it with lysates from HEK293T cells expressing HA-tagged GATOR1 subunits. GST alone and GST-Rac were included as negative controls to account for nonspecific pulldown. We found that only nucleotide-free Rheb robustly pulled down Nprl2 and Nprl3, while neither GDP- nor GTPγS-bound Rheb showed appreciable interaction (**Fig. 4C**). Consistent with our structural predictions, Depdc5 was not recovered in the pulldown, supporting the conclusion that Rheb interacts specifically with the Nprl2/Nprl3 heterodimer, and that this interaction is favored in the nucleotide-free state. Moreover, because the interaction is observed using purified recombinant Rheb incubated with cell lysates and is absent with GDP- or GTPγS-loaded Rheb as well as with GST alone, these results further indicate that the Rheb-Nprl2/3 interaction is biochemically specific and not an artifact of overexpression.

## DISCUSSION

Our study identifies a previously unrecognized point of intersection between the two canonical upstream branches of the mTORC1 pathway, growth factor-regulated Rheb and nutrient-regulated GATOR1. By combining *in vivo* quantitative proteomics in *C. elegans* and biochemical reconstitution in human cells, we demonstrate that inactive, nucleotide-free Rheb associates with the Nprl2/3 subcomplex of GATOR1. This interaction occurs across evolutionary distance, as the association is detectable in *C. elegans*, which lacks the TSC complex, and is conserved in human cells where TSC is present. These findings redefine how cells may coordinate nutrient and growth-factor inputs upstream of mTORC1 and reveal a previously unappreciated layer of regulation in Rheb biology.

A central feature of our findings is the nucleotide-state dependence of this interaction. Only the nucleotide-free conformation of Rheb binds Nprl2/3, whereas GTP-loaded Rheb shows markedly reduced association. This preference has important mechanistic implications. Rheb, unlike Ras or other related GTPases, possesses exceptionally low intrinsic GTPase activity and thus predominantly exists in its GTP-bound form under physiological conditions^30^. This biochemical property, together with the absence of any known Rheb-directed GEF, likely explains why a transient apo-Rheb–binding partner has not been identified previously, because under most cellular conditions the nucleotide-free window for Rheb is extremely short. Only when Rheb is prevented from immediately reloading GTP (e.g., during nutrient stress or in the S20N mutant) does the nucleotide-free conformation accumulate sufficiently to reveal its binding partners. Our data therefore suggest that Nprl2/3 preferentially recognizes a normally rare conformer of Rheb, one that has remained largely inaccessible in prior studies.

Another important conceptual advance from this work is the suggestion that Nprl2/3 may exist and function outside the full GATOR1 complex. This idea is supported by our biochemical and *in silico* findings and is consistent with published data indicating that Nprl2 and Nprl3 can operate outside the full GATOR1 complex. We find that Rheb interacts efficiently with the Nprl2/3 heterodimer, and AlphaFold3 consistently places Rheb on the same region of Nprl3 regardless of whether Depdc5 is included. In contrast, the predicted positioning of RagA is strongly dependent on complex completeness. When RagA is modeled only with Nprl2/3, AlphaFold3 places RagA at the Nprl2 longin domain corresponding to the canonical GAP pocket, but when the complete GATOR1 complex is provided, AlphaFold3 instead positions RagA at the inhibitory Depdc5-binding site as observed in cryo-EM structures^34^. Rheb’s consistent Nprl3 binding, in contrast to RagA’s Depdc5-dependent repositioning, reveals an important mechanistic difference in how Nprl2/3 and the GATOR1 complex regulate the two GTPases. These findings are consistent with emerging evidence that Nprl2 and Nprl3 can function outside the canonical GATOR1 trimer. For example, overexpression of Nprl2 induces NOX2-dependent ROS production, mitochondrial dysfunction, and apoptosis, and proteomic analysis of Nprl2-overexpressing cells identified multiple mitochondrial-associated proteins involved in oxidative stress, electron transport, and apoptotic pathways^36^. Notably, these phenotypes and interactors were observed in the absence of the full GATOR1 complex, indicating that Nprl2 can engage functions independent of Depdc5. Likewise, work in Drosophila emphasized Nprl2 and Nprl3 as a functional unit that inhibits TORC1 during amino acid starvation, highlighting an Nprl2/3-centered module whose behavior is not always explicitly dependent on Depdc5^37^. These observations together with our discovery that inactive Rheb associates specifically with Nprl2/3, but not Depdc5, is consistent with this emerging model that Nprl2/3 forms physiologically relevant subcomplexes with functions that diverge from canonical Rag regulation.

Our findings also raise the possibility that the nucleotide-free state of Rheb may serve regulatory functions under specific physiological conditions. Although transient under steady-state conditions, nucleotide-free intermediates of small GTPases are well documented in other systems and can be selectively stabilized by interacting partners, oxidative stress, or specialized regulatory proteins^38–41^. Given that mTORC1 is acutely responsive to nutrient stress, this raises the question of whether the Nprl2/3–Rheb interaction might serve as a buffering or scaffolding mechanism that prevents premature Rheb activation until favorable metabolic conditions are restored. While our data establish the interaction and define its biochemical determinants, further studies will be required to determine whether Nprl2/3 regulates Rheb localization, stability, or availability to mTORC1 under physiological conditions.

The identification of a physical association between Rheb and Nprl2/3 has important implications for organisms that lack core components of the canonical AND-gate architecture. Both *C. elegans* and *S. cerevisiae* lack the TSC complex yet retain Rheb, PI3K-Akt signaling, and fully functional mTORC1 outputs. The presence of an alternative Rheb-regulatory mechanism in these organisms may help explain how growth-factor and nutrient cues remain coordinated despite the absence of TSC. Moreover, the conservation of this interaction in human cells, where TSC is present, suggests that Nprl2/3-based regulation may operate in parallel with TSC rather than replacing it, perhaps becoming particularly relevant under conditions where Rheb activation needs to be restrained or delayed.

In summary, this study reveals a conserved, nucleotide-state-dependent interaction between Rheb and the Nprl2/3 subcomplex of GATOR1 and highlights a potential mechanism for signal integration upstream of mTORC1. By uncovering this molecular link between the nutrient- and growth-factor–responsive branches of the pathway, our work provides a new framework for understanding how cells might achieve robust and contextually appropriate control of mTORC1 activity.

## METHODS

### C. elegans strains and husbandry

The N2 Bristol wild-type *C. elegans* strain was obtained from the Caenorhabditis Genetics Center (CGC), funded by the NIH Office of Research Infrastructure Programs (P40 OD010440). The strain WBM1611: *rheb-1*(*wbm98*) III [N2, 3xFLAG::*rheb-1*] was generated by inserting a 3xFLAG epitope at the amino terminus of *rheb-1* at its endogenous locus using CRISPR/Cas9-mediated genome editing. The insertion was confirmed by Sanger sequencing (Genewiz, Waltham, MA) and the strain was outcrossed 6 times prior to use. Worms were maintained on standard nematode growth media (NGM) plates seeded with *Escherichia coli* OP50-1 and incubated at 20 °C. OP50-1 bacteria were cultured overnight in Luria-Bertani (LB) broth at 37 °C, after which 100 μl of liquid culture was seeded onto NGM plates to grow for 2 days at room temperature before adding *C. elegans*. For the proteomics experiment, *C. elegans* were grown at 20 °C on the *E*. *coli* strain HT115.

### Microinjection and CRISPR/Cas9 genome editing

CRISPR-mediated genome editing to generate the *C. elegans* strains was performed using a previously described protocol. Briefly, homology-directed repair templates were generated by PCR using primers that appended at least 35 base pairs of homology to both ends of the insert. *dpy-10* was included as a co-injection marker to facilitate selection of edited progeny. CRISPR injection mixes were prepared with the following composition: 0.375 μl HEPES (200 mM, pH 7.4), 0.25 μl KCl (1 M), 2.5 μl trans-activating CRISPR RNA (tracrRNA, 4 μg/μl; Dharmacon, Lafayette, CO), 0.6 μl *dpy-10* CRISPR RNA (2.6 μg/μl), 0.25 μl *dpy-10* ssODN (500 ng/μl), and a PCR repair template (up to 500 ng/μl final concentration). Nuclease-free water was added to bring the final volume to 8 μl. Prior to injection, 2 μl of purified Cas9 protein (12 μg/μl) was added to the mix, mixed by pipetting, spun at 13,000 rpm for 2 minutes, and incubated at 37 °C for 10 minutes. Injection mixes were then introduced into the germline of day 1 adult hermaphrodites via standard microinjection procedures.

### Sample preparation and immunoprecipitation for C. elegans IP-MS

Approximately 300,000 worms were grown across 80 NGM 10 cm plates, each seeded with 3,750 synchronized animals and 1 ml of 20-fold concentrated *E*. *coli* HT115. Bacteria were prepared by pelleting overnight LB cultures and resuspending in fresh LB to achieve a 20x density. N2 and 3×FLAG::RHEB-1 animals were cultured at 20 °C until day 1 of gravid adulthood. Worms were harvested in M9 buffer and washed three times to remove residual bacteria. The final two washes were performed in lysis buffer supplemented with 10% glycerol (100 mM HEPES pH 7.5, 2 mM MgCl₂, 300 mM NaCl, 5 mM 2-mercaptoethanol, 0.05% NP-40), along with protease (8340, Millipore Sigma, St. Louis, MO) and phosphatase (04906837001, Roche, Basel, Switzerland) inhibitors.

After the final centrifugation, excess liquid was removed and worm pellets were weighed. Pellets were snap-frozen by dispensing ∼20 μl drops into liquid nitrogen using a P200 pipette to form small frozen pearls. Worms were lysed using a CryoMill (Retsch, Haan, Germany) with three 1-minute pulses at 30 Hz in the presence of 5 mm metal beads. Pulverized material equivalent to 1 g of worm mass was homogenized in 3 ml of lysis buffer (L3412 Millipore Sigma; 50 mM Tris-HCl pH 7.4, 150 mM NaCl, 1 mM EDTA, 1% Triton X-100) containing protease and phosphatase inhibitors. Total protein concentration was quantified using the Pierce BCA Protein Assay Kit (PI23227, Thermo Fisher, Waltham, Massachusetts) per manufacturer’s instructions.

Anti-FLAG M2 magnetic beads (M8823, Millipore Sigma) were washed three times in 1× wash buffer (W0390 Millipore Sigma; 0.5 M Tris-HCl pH 7.4, 1.5 M NaCl) and then incubated with worm lysates overnight at 4 °C with end-over-end rotation. After binding, beads were washed three times with 1 ml of wash buffer containing protease and phosphatase inhibitors, using 2-minute end-over-end rotation followed by 2 minutes on a magnetic stand. Proteins were eluted using 3×FLAG peptide (F4799, Millipore Sigma) at a final concentration of 300 ng/μl, prepared by dissolving 1 mg of peptide in 40 μl of 10× wash buffer followed by 160 μl distilled water. Beads were incubated with 125 μl of elution buffer for 4 h at 4 °C. Eluates were collected by magnetic separation and transferred to clean tubes. A 5% aliquot of the eluate was analyzed by SDS-PAGE and silver staining for quality control, and the remaining sample was used for mass spectrometry. Four independent biological replicates were processed using this workflow.

### MS acquisition and analysis pipeline

Mass spectrometry was performed at the Thermo Fisher Center for Multiplexed Proteomics (TCMP) using an Orbitrap Eclipse™ Tribrid™ mass spectrometer and isobaric tandem mass tags (TMT) for quantitative proteomics^17^. Following immunoprecipitation, protein concentration was estimated, and samples were processed in a washing buffer containing 1 M urea and 50 mM Tris (pH 8.0). Proteins were reduced with 5 mM TCEP in wash buffer, alkylated with 5 mM iodoacetamide, and the reaction quenched with 10 mM DTT. Precipitation was performed using trichloroacetic acid (TCA), and pellets were resuspended and digested with trypsin in 1 M urea, 50 mM Tris (pH 8.0). Resulting peptides were labeled with TMT reagents, and ∼2 μl of each labeled sample was combined to confirm labeling efficiency. Ratio checks indicated 99.6% TMT label incorporation. Labeled peptides were pooled into a single multiplexed sample, desalted using stage tips, and loaded onto an in-house packed capillary column for liquid chromatography separation based on peptide hydrophobicity and charge. Eluted peptides were ionized and analyzed by mass spectrometry (LC-MS) on the Orbitrap Eclipse platform. MS spectra were searched using a Sequest-based in-house software platform with the following parameters: peptide mass tolerance, 50 ppm; fragment ion tolerance, 1.0 Da; maximum internal cleavage sites, 2; and maximum differential modification sites, 4. MS2 spectra were searched using the SEQUEST algorithm against a *C. elegans* UniProt composite database containing reversed sequences and known contaminants. Peptide spectral matches were filtered to a 1% false discovery rate (FDR) using the target-decoy strategy combined with linear discriminant analysis. Quantification was performed using peptides with a summed signal-to-noise (S/N) threshold > 80 and isolation specificity (IS) > 0.5. All peptides (unique and redundant) were matched and aggregated into protein groups. A total of 13,627 peptides were identified and assembled into 2,495 protein groups. After filtering, 2,405 proteins met the quantification criteria and were included in the final dataset.

### Mammalian cell culture and transfections

HEK293T control, TSC2^−/−^, and HA-Depdc5 cell lines were maintained in DMEM supplemented with 10% FBS and 1% penicillin-streptomycin in a humidified incubator at 37°C with 5% CO₂. Transient transfections were performed using Lipofectamine™ 3000 Transfection Reagent (L3000015, Thermo Fisher Scientific, Waltham, MA). For co-immunoprecipitation (CO-IP) experiments, 0.3-0.5 million HEK293T cells were seeded in 6-well cell culture plates. 24 h later, cells were transfected with combinations of the following plasmids: HA-GST-Rheb, FLAG-Rheb, FLAG-Rheb^S20N^, FLAG-Rheb^Q64L^, and FLAG-Rheb^C181S^. Additional plasmids included pRK5 HA-RagA (#99710), pRK5 HA-Nprl2 (#99709), pRK5 FLAG-Nprl2 (#46333), and pRK5 HA-Nprl3 (#46330), all obtained from Addgene (Watertown, MA). After 36-48 h post-transfection, cells were washed twice with PBS and subjected to the indicated nutrient or pharmacological treatments. For mTORC1 pathway inhibition, cells were treated with vehicle (DMSO), 1 μM wortmannin, 100nM rapamycin (R-5000, LC Laboratories, Woburn, MA) or 250 nM torin (4247, Tocris Bioscience, Bristol, United Kingdom) for 1 h. For serum starvation, cells were incubated in DMEM lacking FBS for 16 h. For amino acid starvation, cells were preconditioned in RPMI medium containing amino acids and 10% FBS for 16 h, washed twice with PBS, and then transferred to RPMI supplemented with 10% dialyzed FBS and lacking amino acids and glucose for 4 h.

### Co-immunoprecipitation

Co-immunoprecipitation and cell signaling experiments were performed based on established protocols^34,42^. Cells were rinsed with ice-cold PBS and lysed in Triton buffer (1% Triton X-100, 40 mM HEPES pH 7.4, 2.5 mM MgCl₂, 100 mM NaCl) supplemented with protease and phosphatase inhibitors. Lysates were clarified by centrifugation, and the supernatant (soluble fraction) was transferred to a fresh tube for protein quantification using the Bicinchoninic Acid (BCA) assay (23227, Thermo Fisher Scientific) according to the manufacturer’s instructions. Equal amounts of clarified lysate were then incubated with anti-FLAG® M2 magnetic beads (M8823, Millipore Sigma) overnight at 4 °C. Beads were washed four times in lysis buffer and proteins eluted by boiling in SDS sample buffer. Samples were resolved by SDS-PAGE and analyzed by immunoblotting.

### Immunoblotting

After transfer, membranes were segmented at defined molecular weight regions based on the protein ladder to allow probing with multiple primary antibodies. The following rabbit monoclonal antibodies (Cell Signaling Technology, Danvers, MA) were used at 1:1,000 dilution: anti-HA (C29F4, #3724), anti-RagA (D8B5, #4357), anti-β-Actin (#4967), anti-phospho-p70 S6 kinase (Thr389) (108D2, #9234), anti-p70 S6 kinase (49D7, #2708), anti-Tuberin/TSC2 (D93F12, #4308), and anti-FLAG (#14793). Mouse monoclonal anti-FLAG® M2 antibody (Millipore Sigma, #F1804) was used at 1:2,000 dilution. Secondary antibodies included HRP-conjugated anti-mouse IgG (1:5,000, CST #7076) and anti-rabbit IgG (1:2,000, CST #7074). Membranes probed with Cell Signaling Technology antibodies were blocked and incubated in 1× TBST (10 mM Tris-HCl pH 8.0, 150 mM NaCl, 0.05% Tween-20) containing 5% bovine serum albumin (BSA). For all other immunoblots, including those probed with secondary antibodies, 5% non-fat dry milk in TBST was used. Primary antibody incubations were carried out at 4 °C for 16 h and secondary antibody incubations were carried out at room temperature for 1 h. Signal detection was performed by chemiluminescence using a Bio-Rad ChemiDoc XRS+ system (1708265, Bio-Rad, Hercules, CA). Band intensities were quantified using Image Lab Software (Bio-Rad) and normalized to input and loading controls.

### GST pulldown assays

Recombinant GST, GST-Rac1, or GST-Rheb proteins were expressed in *E*. *coli* BL21 cells, with expression induced for 6 hours at 30 °C once cultures reached an optical density (OD₆₀₀) of 0.6-0.8. Cell pellets from 1 L culture were resuspended in 20 ml Rheb lysis buffer (50 mM HEPES, 140 mM NaCl, 1 mM EDTA, 1 mM DTT, and 1:100 protease inhibitor cocktail), followed by sequential treatments with 0.25 mg/mL lysozyme for 15 minutes, 0.2% Triton X-100 for 30 minutes, and 2-5 units of Benzonase nuclease with 12 mM MgCl₂ for 30 minutes. During each step of lysis, samples were gently rotated end-over-end at 4 °C. Lysates were cleared by centrifugation at 20,000 × g, snap-frozen in liquid nitrogen, and stored at −80 °C for long-term preservation.

For pulldown assays, HA-Depdc5 stable cell line co-transfected with HA-Nprl2 and HA-Nprl3 were lysed in ice-cold CHAPS lysis buffer (40 mM HEPES pH 7.4, 120 mM NaCl, 1 mM EDTA, 0.3% CHAPS, 5% glycerol, 10 mM sodium pyrophosphate, 10 mM β-glycerophosphate) supplemented with 1 μM Microcystin-LR (Enzo Life Sciences, ALX-350-012-C500) and 1:100 protease inhibitor cocktail (Sigma, P8340). Lysates were clarified by centrifugation and total protein concentration was normalized across samples using the BCA assay. Recombinant GST-tagged proteins were pre-bound to glutathione agarose beads (Pierce, PI16101) and washed three times in Rheb wash buffer (50 mM HEPES pH 7.4, 500 mM NaCl, 2 mM EDTA, 0.2% Triton X-100, 1 mM DTT, and 1:100 protease inhibitors), followed by two washes in nucleotide loading buffer (50 mM HEPES pH 7.4, 1 mM EDTA, 5 mg/mL BSA, and 1:200 protease inhibitors). Beads were incubated with 500 μM GDP or GTPγS in loading buffer for 15 minutes at 30 °C with gentle agitation, then placed on ice and incubated with 10 mM MgCl₂ for 5 minutes to lock in nucleotide binding.

Normalized cell lysates (1 mL at 2 mg/ml concentration) were added to 20 μL of GST-bound glutathione bead slurry (1:1 bead:buffer). For nucleotide-free conditions, 5 mM EDTA was added. For GDP- or GTPγS-bound conditions, 10 mM MgCl₂ and 500 μM of the corresponding nucleotide were supplemented. Pulldown reactions were incubated at 4 °C for 4 hours with end-over-end rotation. Beads were washed four times with CHAPS lysis buffer containing 10 mM MgCl₂ and proteins were eluted by boiling in SDS sample buffer at 70 °C for 10 minutes prior to SDS-PAGE and immunoblot analysis.

### AlphaFold3 modeling

All structural predictions were performed using AlphaFold3^15^ on the AlphaFold server (https://alphafoldserver.com/). Full-length sequences of human Rheb (WT, S20N, Q64L), Nprl2, Nprl3, Depdc5, or RagA were input. For each condition, models were generated using 2 seeds with five models per seed (total 10 models per condition). The best-ranked model was selected for per-residue pLDDT scores, inter-subunit interface predicted TM-scores (ipTM), and global pTM scores. These score were extracted from the summary_confidences_X.json files. PAE matrices were retrieved from full_data_X.json. Ribbon structures were rendered using ChimeraX with pLDDT values mapped to B-factor color gradients.

### Statistical analyses

Statistical analyses were performed in R studio (Posit Software, Boston, MA) and GraphPad Prism 10 (GraphPad Software, La Jolla, CA) using the statistical tests indicated in figure legends.

Predicted Aligned Error (PAE) matrices for each AlphaFold3 model were extracted from the full_data_X.json files generated by the Alphafold server. For each model, the *full_data_X.json* file was used to extract the full N×N PAE matrix and corresponding chain and residue token indices. Custom Python scripts were used to calculate interface-level confidence metrics based on PAE values between contacting residues. Model coordinates were parsed from the associated *.cif* structure files and loaded using the Biopython MMCIFParser module. Contacting residues between chains were defined as residue pairs with a Cα-Cα distance ≤ 8 Å. Median PAE was calculated symmetrically across the interface using only residue pairs that met the distance cutoff in either direction (i.e., A-to-B or B-to-A). For each condition (WT, S20N, and Q64L), models from all ranked outputs (typically 5 per AlphaFold run) and multiple seeds (when used) were analyzed. Models without any residue pairs under the 8 Å threshold were classified as “no contact” cases and excluded from the contact PAE average but tallied separately to assess reproducibility. All calculations were performed using NumPy and Pandas, and plots were generated using GraphPad Prism. Custom scripts used for PAE extraction and filtering are available upon request.

## ACKNOWLEDGEMENTS

HEK293T stably expressing Depdc5-HA were generously provided by Dr. Kuang Shen (UMass Chan Medical School). HEK293T TSC2^−/−^ cell line and bacterial and mammalian Rheb plasmids were kindly provided by Dr. Brendan Manning (Harvard T.H. Chan School of Public Health). Shin Jong and Sophie Lockwood Evarts helped with experiments. We thank Julian Mintseris and Jon Van Vranken from the Thermo Fisher Center for Multiplexed Proteomics at Harvard Medical School for the IP-mass spectrometry. We are also grateful to members of the Mair laboratory for helpful discussions on the project and the manuscript. W.B.M. is funded by NIH/NIA R01AG067106 and R01AG044346.

## AUTHOR CONTRIBUTIONS

A.P and W.B.M conceived the idea for the study. A.P performed the experiments and analyzed the data. A.P and W.B.M wrote the manuscript.

## SUPPLEMENTAL FIGURE LEGENDS

**Figure S1.**
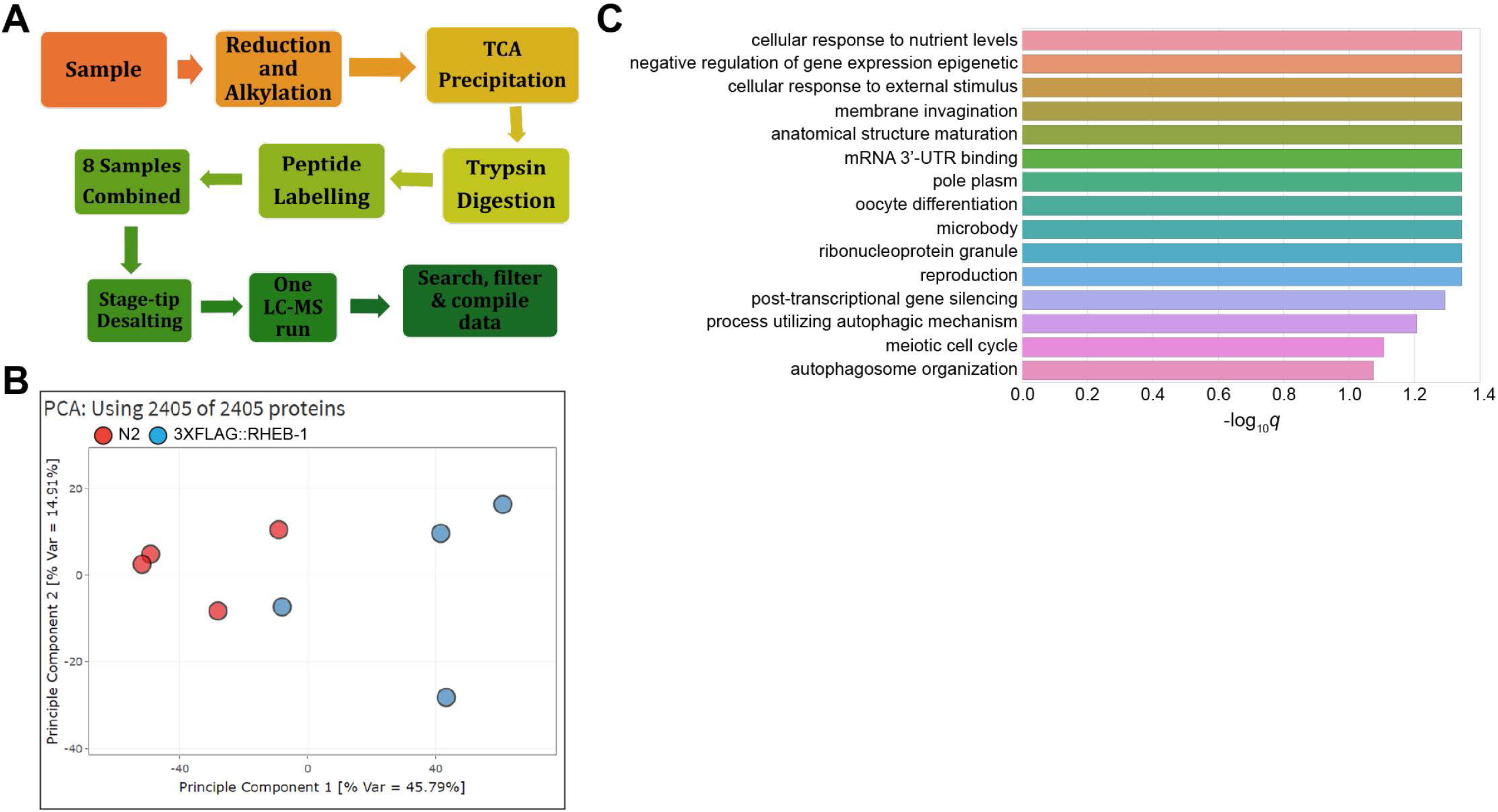
Identifying *C. elegans* RHEB-1 interactors via TMT-based quantitative proteomics. **(A)** Schematic outlining the tandem mass tag (TMT)-based quantitative proteomics pipeline. Following FLAG immunoprecipitation from *C. elegans* lysates, protein samples were quantified, reduced, alkylated, and TCA-precipitated before enzymatic digestion with trypsin. Peptides were labeled with TMT reagents and pooled into a single multiplexed sample. After desalting, peptides were separated by liquid chromatography and analyzed by mass spectrometry (LC-MS). MS2 spectra were searched against a *C. elegans* UniProt database using a Sequest-based in-house platform. Peptides were filtered to a 1% false discovery rate (FDR) and proteins were quantified based on summed signal-to-noise (S/N) > 80 and isolation specificity (IS) > 0.5. In total, 13,627 peptides were mapped to 2,495 protein groups, with 2,405 proteins meeting quantification criteria. **(B)** Principal component analysis (PCA) of the final dataset using all 2,405 quantified proteins across four biological replicates each from 3×FLAG::RHEB-1 and control samples. The clear separation between conditions reflects high data quality and reproducibility. **(C)** Gene Ontology (GO) enrichment analysis of the top 90 RHEB-1-associated proteins identified by TMT proteomics. These hits passed both a log₂ fold-change cutoff of >2 and a p-value threshold of <0.01. Enrichment was performed using WormBase gene set annotations.

**Figure S2.**
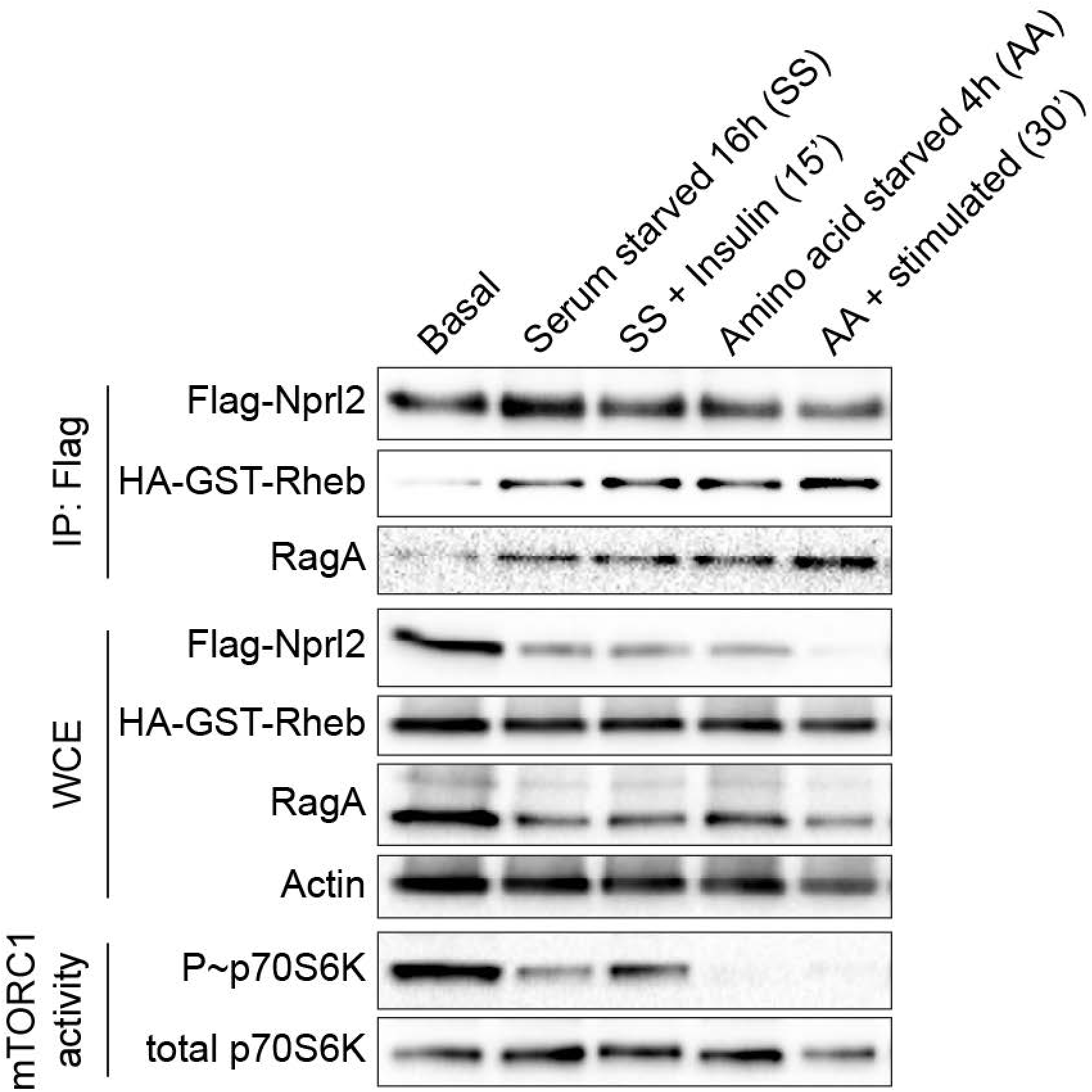
Serum and amino-acid starvation enhance the Rheb-Nprl2 interaction in HEK293T cells. HEK293T cells expressing HA-GST-Rheb and FLAG-Nprl2 were treated as indicated. Co-immunoprecipitation was performed on the lysates. Total expression shown in whole-cell extracts (WCE). mTORC1 activity was monitored by phospho-p70S6K levels in whole-cell extracts. RagA was included as a positive control.

**Figure S3.**
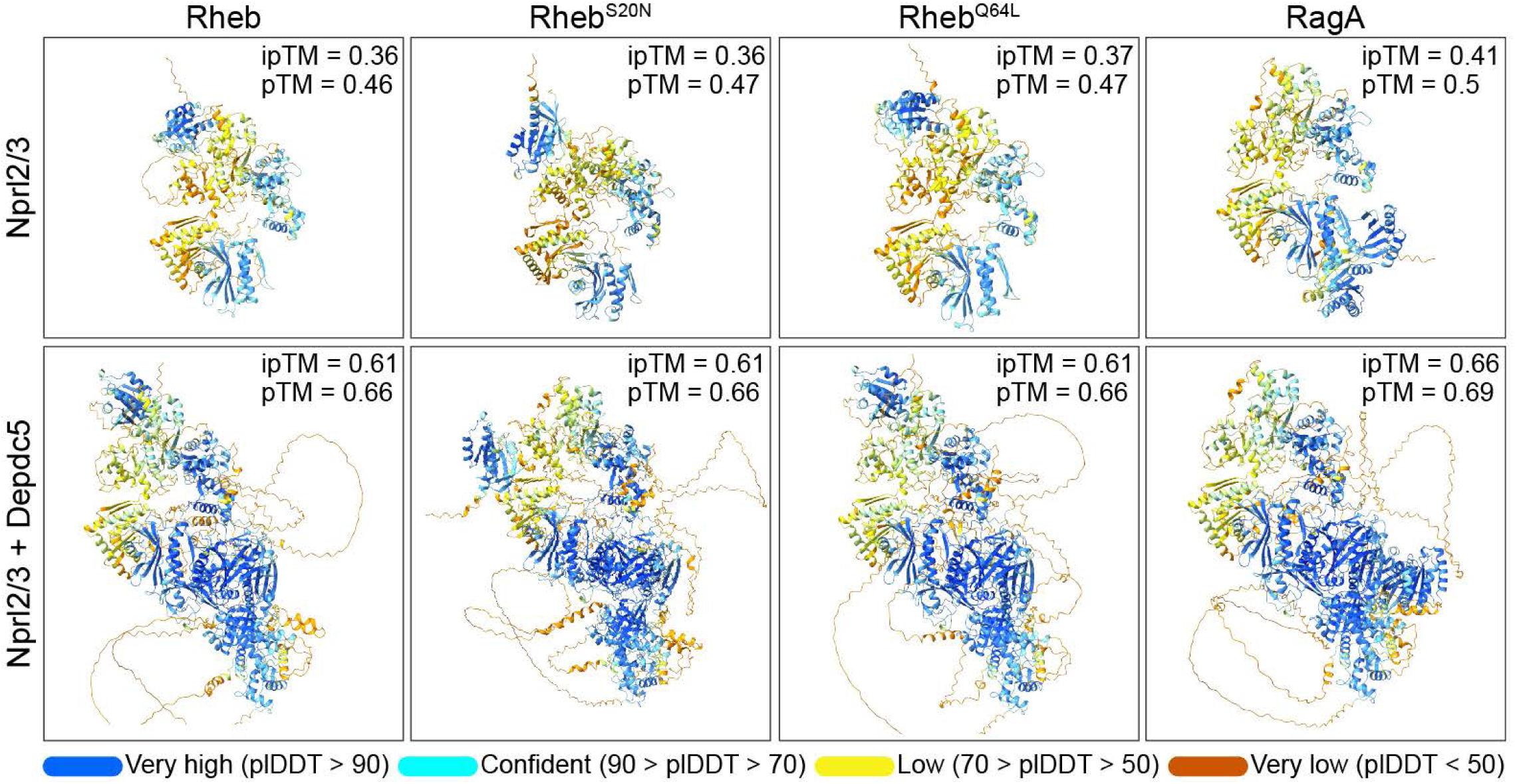
AlphaFold3 model confidences for Rheb- and RagA-GATOR1 complexes. AlphaFold3-predicted structures of human Rheb (wild-type, S20N, Q64L) or RagA modeled with Nprl2/3 alone (top) or with the full GATOR1 complex (Nprl2/3 + Depdc5; bottom). Color corresponds to predicted per-residue confidence (pLDDT), ranging from very low (orange) to very high (blue). Models lacking Depdc5 exhibit lower overall confidence and increased disorder across the complex, reflected in both reduced pLDDT and lower ipTM and pTM scores. Inclusion of Depdc5 markedly improves global fold quality (pTM) and inter-subunit interface confidence (ipTM), even though Depdc5 does not directly contact Rheb in these models.

## Notes

### Competing Interest Statement

The authors have declared no competing interest.

